# Reversed paired-gRNA plasmid cloning strategy for efficient genome editing in *Escherichia coli*

**DOI:** 10.1101/839555

**Authors:** Tingting Ding, Chaoyong Huang, Zeyu Liang, Xiaoyan Ma, Ning Wang, Yi-Xin Huo

## Abstract

A growing number of CRISPR-Cas9 associated applications require co-expression of two distinct gRNAs. However, coexpressing paired gRNAs under the driving of independent but identical promoters in the same direction triggers plasmid instability, due to the presence of direct repeats (DRs). In this study, deletion between DRs occurred with high frequencies during plasmid construction and duplication processes, when three DRs-involved paired-gRNA plasmids cloning strategies were tested. This recombination phenomenon was RecA-independent, in agreement with the replication slippage model. To completely eliminate the DRs-induced plasmid instability, a reversed paired-gRNA plasmids (RPGPs) cloning strategy was developed by converting DRs to the more stable invert repeats (IRs). Using RPGPs, we achieved a rapid deletion of chromosome fragments up to 100 kb with high efficiency of 83.33% in *Escherichia coli*. This study provides general solutions to construct stable plasmids containing short DRs, which can improve the performances of CRISPR systems that relied on paired gRNAs, and also facilitate other applications involving repeated genetic parts.

## Introduction

CRISPR-based systems are powerful tools for genetic manipulations in both eukaryotic and prokaryotic organisms, which solely relies on single guide RNA molecule (gRNA) for targeting (Jinek *et al.*, 2012; Jiang *et al.*, 2013). CRISPR applications have been usefully extended when two distinct genomic locations targeted simultaneously. For example, paired gRNAs are required to dramatically reduce off-target mutations (Ran *et al.*, 2013), to achieve combinatorial genome modifications (Li *et al.*, 2015) and to facilitate large genomic deletion (Su *et al.*, 2016; Zheng *et al.*, 2017). The stable co-expression of paired gRNA determines the precision, the reliability, and the efficiency of CRISPR applications.

Although various cloning strategies have been established for the expression of paired or multiple gRNAs, the instability of gRNA plasmids is still an urgent problem to be solved. In general, multiple targeting spacers can be expressed by the CRISPR array or gRNA cassettes. However, plasmids with direct repeats (DRs) are difficult to assemble *in vitro* and lead to genetic instability *in vivo*. When using CRISPR array, recombination between DRs of the array causes the targeting spacer sequence eliminated and then genomic modification failed (Jiang *et al.*, 2013; Su *et al.*, 2016). Similarly, coexpressing multiple gRNAs in one plasmid results in self-homologous recombination if gRNA cassettes are transcribed by independent but identical promoters in the same direction (Aparicioprat *et al.*, 2015; Jiang *et al.*, 2015; Vidigal and Ventura, 2015). To eliminate recombination, the building of promoter library and functional gRNA scaffold library (Reis *et al.*, 2019) can be further explored, but it may be limited by optional constructive promoters and complex machine learning technology. Thus, we attempt to investigate the recombination mechanism of DRs-involved paired-gRNA plasmids and develop simple but universal cloning strategies to prevent DRs-mediated recombination genetically.

Normally, the rearrangement of DRs are induced by recombinational or replicational mechanism in *E. coli* (Lovett *et al.*, 1993). The former is mediated by recombinase A (RecA) which promotes the pairing and the strand exchange between homologous sequences to form heteroduplex DNA (Cox and Lehman, 1987; Radding, 1989; Kowalczykowski, 1991). For repeats larger than about 200 bp in length, deletion can occur by RecA-dependent homologous recombination (Bi and Liu, 1994). For homologies less than about 200 bp in length, the plasmid rearrangement can occur in a RecA-independent manner (Bi and Liu, 1994; Bi *et al.*, 1995; Bi and Liu, 1996a; Azpiroz and Laviña, 2017). Depending on plasmid recombination products and the inverting sequences between DRs, three mechanisms of RecA-independent rearrangements were presented: slipped misalignment, sister-chromosome slipped misalignment, and single-strand annealing (Bzymek and Lovett, 2001). Thus, the recombination mechanism of DRs-involved paired-gRNA plasmids can be predicted if tested in organisms with different *recA* genotype.

In this study, we investigated the deletion rates and deletion types of three DRs-involved paired-gRNA plasmids cloning strategies and showed a replication mechanism for gRNA deletion. All of these plasmids could not thoroughly eliminate recombination during assembly and re-transformation processes, as long as DRs existed. Thus, a simple reversed paired-gRNA plasmids (RPGPs) cloning strategy was developed by placing paired gRNA cassettes in the opposite direction, which converted DRs to invert repeats (IRs). RPGPs can thoroughly avoid DRs-mediated recombination during DNA replication. Thus, they have a great potential to facilitate the construction of large-scale gRNA libraries for CRISPR applications that require co-expression of two gRNAs. Our strategy can also be applied for the construction of other plasmids containing repeated genetic parts.

## Results

### The stability of DRs-involved paired-gRNA plasmids pDG-A-X series

To study the stability of DRs-involved paired-gRNA plasmids, pDG-A-X series for co-expression of two gRNAs were employed. A functional gRNA contains a 20-bp sequence for targeting and a 82-bp scaffold that binds Cas9 protein (Li *et al.*, 2015). Each gRNA was transcribed by a 35 bp constitutive promoters J23119 **(Figure 1A)**. Plasmid rearrangement was detected by PCR primers F1/R1 after plasmid construction and re-transformation processes.

**Figure 1.**
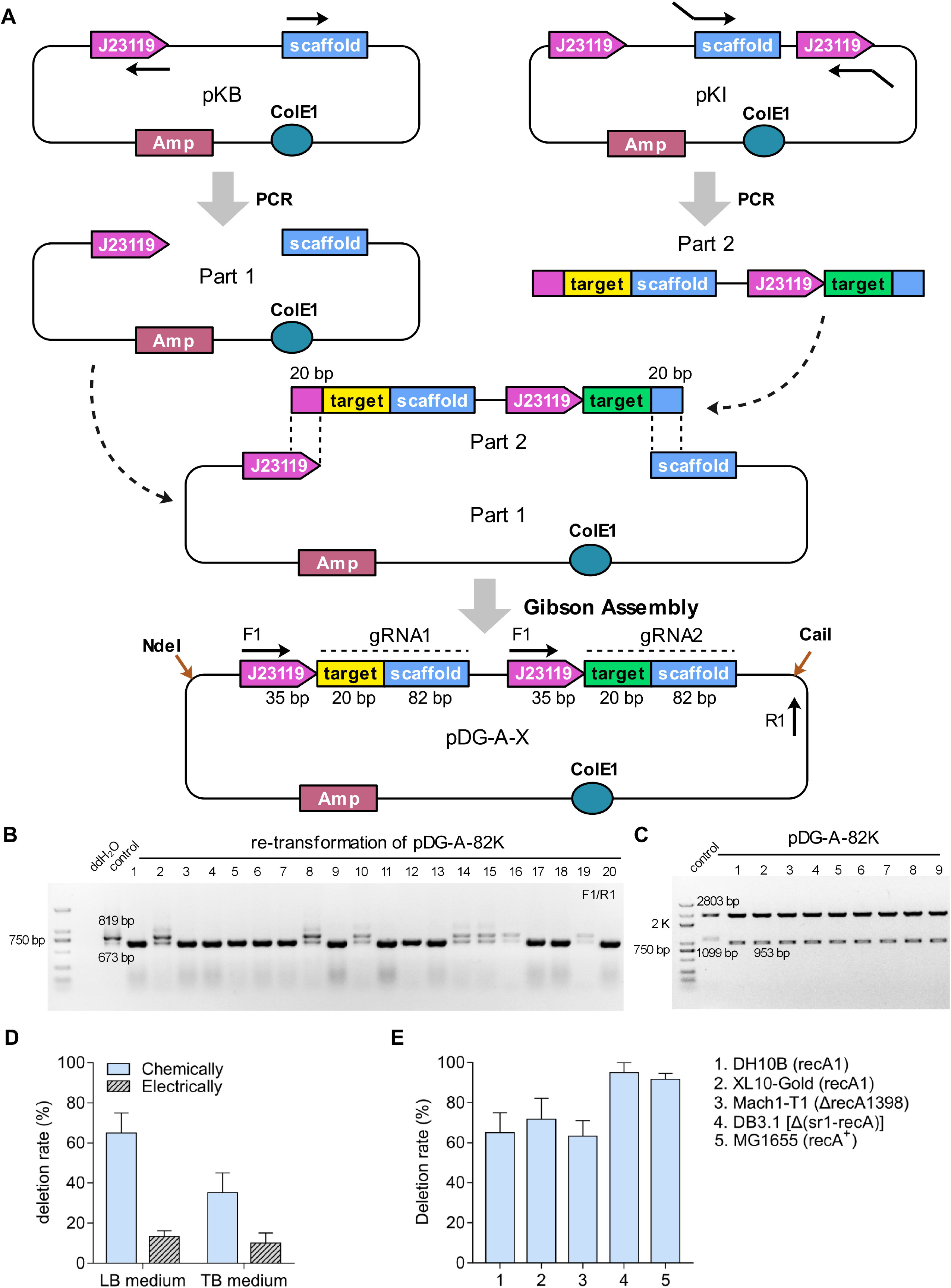
The design and stability of DRs-involved paired-gRNA plasmids pDG-A-100K in *E. coli*. A) The modular construction strategy of pDG-A-X series. pKB plasmid was used for PCR amplification of DNA part 1, which contained pDG-A-X backbone, one constitutive promoter J23119, and a gRNA scaffold. pKI plasmid was used for PCR amplification of part 2 series, which contained a gRNA fragment followed by another constitutive promoter J23119 and 20-bp target sequence. For the PCR reaction, the 20-bp sequences specific for two targeted loci and the 20-bp overlap sequences for Gibson Assembly were embedded in primers as a part of insert. Gibson Assembly method was preformed to assemble these parts into pDG-A-X series. B) Representative PCR results for the deletion rate of pDG-A-100K after re-transformation process in *E. coli*. C) The double restriction enzyme digestion analyses of pDG-A-100K and its deletion derivations. D) The deletion rates of pDG-A-100K when introduced by chemical transformation or electroporation and cultured in the condition of Luria-Bertani (LB) medium or Terrific Broth (TB) medium. Data are expressed as means ±s.d. from three independent experiments. E) The deletion rates of pDG-A-100K after the re-transformation process in various strains of *E. coli*. Data are expressed as means ±s.d. from three independent experiments.

pDG-A-100K for 100-kb genomic deletion was constructed by using *E. coli* DH10B strain as host. However, the deletion rate of 73.33% was observed after pDG-A-100K plasmid construction process. Similarly, the deletion rate was around 65% after re-transformation process of the correct pDG-A-100K plasmid **(Figure 1B and 4B)**. PCR results indicated that the deletion occurred between the paired-gRNA regions of these mutant plasmids. DNA sequencing results demonstrated one of two gRNAs with its promoter was eliminated. Furthermore, the double restriction enzyme digestion analyses by using NdeI and CaiI showed the deletion only occurred between the paired-gRNA regions, rather than other parts of plasmids (**Figure 1C**).

To enhance the stability of pDG-A-100K, the effects of experimental conditions including DNA transformation methods and culture mediums were then assessed during re-transformation process in DH10B strain. Compared with transformation by heat shock, electrotransformation led to a 5.6-fold decrease in the deletion rate for cells cultured in LB medium and a 3.5-fold decrease for cells cultured in TB medium **(Figure 1D)**. The nutrient supplies for plasmid propagation also influenced its stability. Replacing the LB medium with the nutrient-rich TB medium reduced the deletion rate by half when DNA was chemically transformed into cells, while no further decrease in the deletion rate was achieved when plasmids were transformed electrically **(Figure 1D)**. Therefore, pDG-A-100K appeared to be more stable when introduced into cells by electroporation and propagated in rich medium, but neither could eliminate the events of plasmid rearrangement.

### The patterns of plasmid rearrangement

Various recombination derivations were discovered during DRs-mediated recombination events of pDG-A-X series **(Figure 2)**. Based on our observations, two main types of deletion were summarized: the deletion of the first gRNA expression cassette along with its promoter (MUT-1), and the deletion of the second gRNA expression cassette together with its promoter (MUT-2). In addition, point mutations in the –10 regions or –35 regions of promoter J23119 (MUT-3/4) appeared frequently, which could affect the transcription process of gRNAs. A 12-bp repeated insertion at the end of the gRNA scaffold was also detected (MUT-5), which could influence the normal structure of gRNA. Taken together, pDG-A-X series produced the deletion of one of paired gRNA expression cassettes randomly and other spontaneous mutation between the paired-gRNA regions, which made it difficult to maintain plasmid stability.

**Figure 2.**
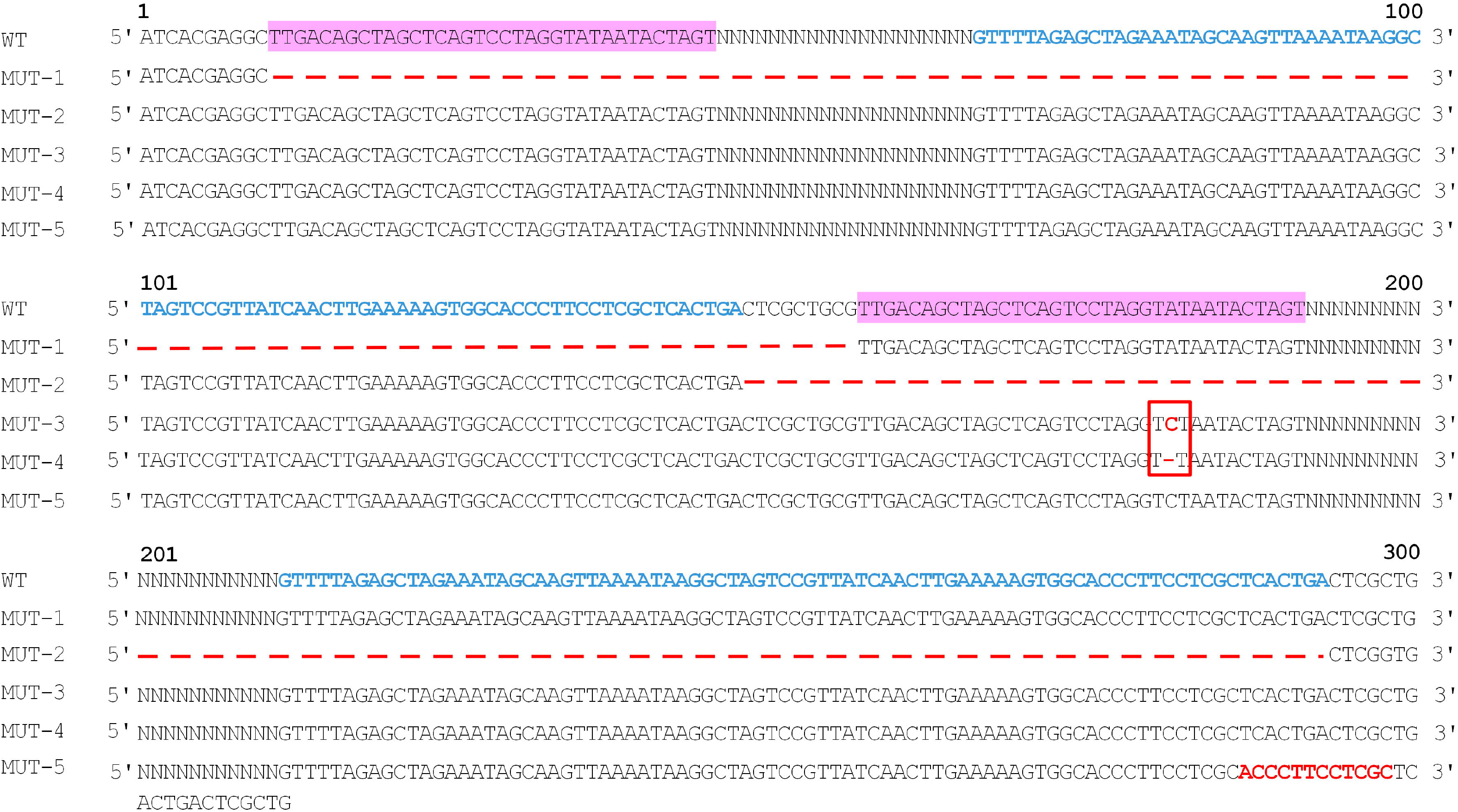
Representative DNA sequencing results of recombination derivatives from pDG-A-X series. MUT-1 had deletion of the first gRNA expression region; MUT-2 had deletion of the second gRNA expression region; MUT-3 and MUT-4 had point mutations of the second promoter J23119; MUT-5 had 12-bp repeated insertion in between the second gRNA scaffold.

### The RecA dependency of paired-gRNA plasmids recombination

To test whether the recombination of pDG-A-X series relied on the RecA enzyme, the correct pDG-A-100K was re-transformed into various *E. coli* strains with the genotypes of *recA1*, Δ*recA1398*, Δ(*sr1-recA*) or *recA*^+^ **(Figure 1E)**. In MG1655 *recA*^+^ strain expressing functional RecA protein, the deletion rate was up to 91.67%. In various *recA* mutant strains, DRs-induced recombination still occurred with the frequencies of 63.33-95%. No distinct difference of deletion rates was found in XL10-Gold *recA*1 strains, DH10B *recA*1 strain and Mach1T1 Δ*recA*1398 strain, while the deletion rate even was increased up to 95% in DB3.1 Δ(*sr1-recA*) strain. All results indicated that RecA-independent recombination played a great role on the deletion of pDG-A-X series in *E. coli*.

The replication slippage model for RecA-independent recombination (Lovett and Feschenko, 1996; Bzymek and Lovett, 2001) was applied to explain the phenomenon, since the main recombination products of pDG-A-X series were plasmid deletion form. The pDG-A-X series have ColE1 origin which produces high plasmid copy numbers and determines unidirectional replication (Del *et al.*, 1998). During the plasmid replication process in *E. coli*, pDG-A-X series generated the Types I or Type II slipped misalignment of the Okazaki fragment, which formed a loop within the lagging strand template to facilitate the formation of deletion **(Figure 3)**. When the second promoter J23119 was employed as mispaired position, the deletion of the first gRNA expression region occurred, leading to the formation of pDG-A-X-M1. When the repeated gRNA scaffold mediated the plasmid recombination, the second gRNA region was deleted to form pDG-A-X-M2.

**Figure 3.**
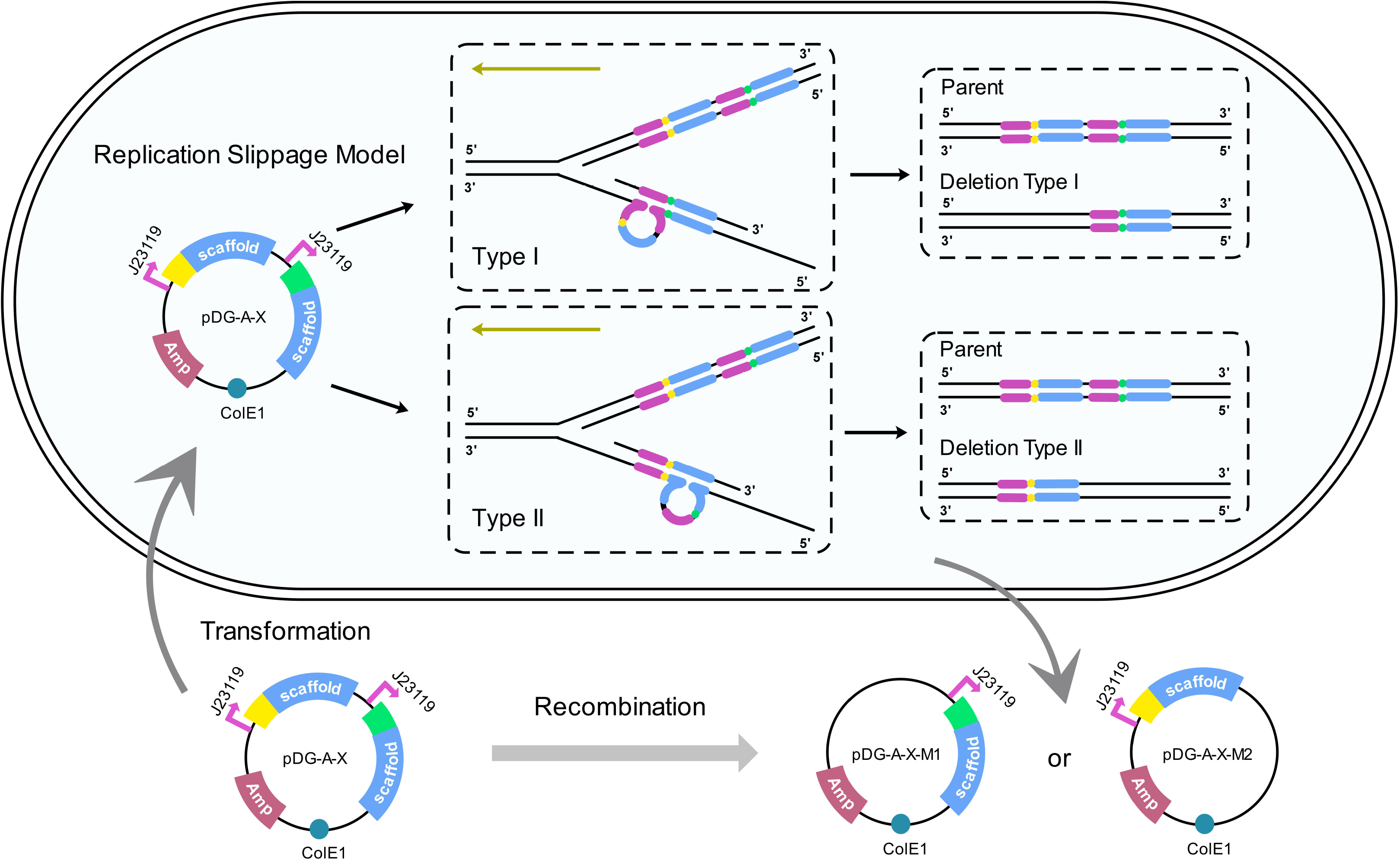
The deletion mechanism of pDG-A-X series during DNA replication. Plasmid pDG-A-X was designed for double gRNA expression: gRNA1 and gRNA2. Each gRNA containing a 20-nt guide sequence (yellow or green) and an 82-bp scaffold (blue) was transcribed by a constitutive promoter J23119 (purple). During the DNA replication process in *E. coli*, pDG-A-X series generated the Types I or Type II slipped misalignment of the Okazaki fragment, which formed a loop within the lagging strand template to facilitate the formation of deletion. Deletion Type I: When the second promoter J23119 was employed as mispaired position, the deletion of the first gRNA expression region occurred, leading to the formation of pDG-A-X-M1. Deletion Type II: When the repeated gRNA scaffold mediated the plasmid recombination, the second gRNA expression region of pDG-A-X series was deleted to form pDG-A-X-M2. The golden arrows indicate the direction of plasmid replication.

### Effects of promoters and origins on pDG-A-X series stability

The effects of different plasmid architecture on plasmid stability were then evaluated. As shown in **Figure 1**, there are two pairs of DRs in pDG-A-X. One is the repeated 35-bp promoter J23119 while the other one is the repeated 82-bp gRNA scaffold. To reduce the number of DRs, pDG-P-X was designed by replacing the second promoter J23119 with an alternative 49-bp PR promoter **(Figure 4A)**. After the assembly products of pDG-P-100K were introduced into DH10B strain, the deletion rate of pDG-P-X was up to 81.67% when verified by primers F1/R1 **(Figure 4B)**. These deletion derivatives of pDG-P-100K didn’t contain promoter PR region when verified by primers F2/R1. DNA sequencing demonstrated that pDG-P-100K generated spontaneous deletion of the second gRNA region to form pDG-A-100K-M2. Although it was difficult to obtain correct pDG-P-X series plasmids by Gibson Assembly method, these plasmids could be more stably maintained during the re-transformation process once the correct plasmid was obtained firstly **(Figure 4B)**.

**Figure 4.**
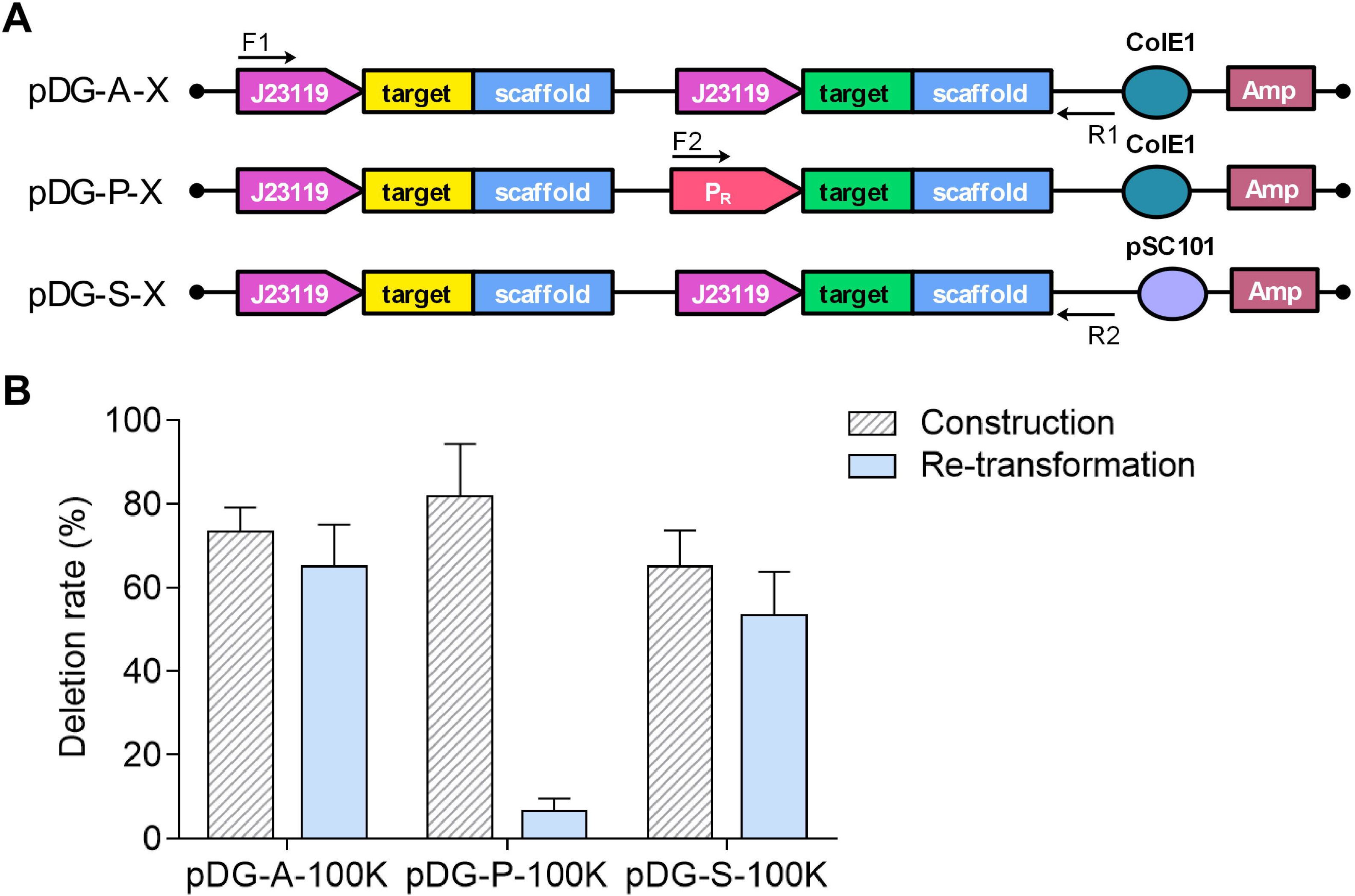
The comparisons of stability of DRs-involved paired-gRNA plasmids pDG-A-X, pDG-P-X and pDG-S-X series. A) Diagrams of pDG-A-X, pDG-P-X and pDG-S-X series. B) The deletion rates of pDG-A-X, pDG-P-100K and pDG-S-100K during plasmid construction and re-transformation processes in DH10B strain. Data are expressed as means ±s.d. from three independent experiments.

Since the copy number may have an impact on plasmid stability, pDG-S-X series were designed by replacing the ColE1 origin with pSC101 origin (Hasunuma and Sekiguchi, 1977) which replicates at a relatively low copy number (<8 copies/cell) **(Figure 4A)**. However, pDG-S-100K still had a high deletion rate of 65% and 53.33% during plasmid construction and re-transformation processes **(Figure 4B)**. These results demonstrated that just changing the promoter or the origin of DRs-involved paired-gRNA plasmids pDG-A-X series didn’t eliminate the events of plasmid rearrangement.

### Design of reversed paired-gRNA plasmids (RPGPs) cloning strategy

In attempt to avoid DRs-induced plasmid rearrangement genetically, a reversed paired-gRNA plasmids (RPGPs) cloning strategy was developed for pDG-R-X series **(Figure 5A)**. Compared with pDG-A-X, the plasmid architecture of pDG-R-X was versatilely modified through changing the promoter of the second gRNA, the origin of replication, and the direction of gRNA cassettes. Two gRNA cassettes were placed in opposite directions with one expressed by J23119 promoter and another by P_R_ promoter, thus turning the two 82-bp gRNA scaffolds into inverted repeats (IRs). During plasmid construction process, the 20-bp sequences specific for two targeted loci and the 20-bp overlap sequences for assembly were embedded in primers as a part of insert. Since the overlap sequences were repeated but reversed, the insert could be assembled in two directions, leading to the formation of pDG-R1-X or pDG-R2-X **(Figure 5A)**. As we expected, pDG-R-100K didn’t generate any plasmid rearrangement events during plasmid construction process, when verified by PCR reactions (F3/R3 and F4/R2) and DNA sequencing.

**Figure 5.**
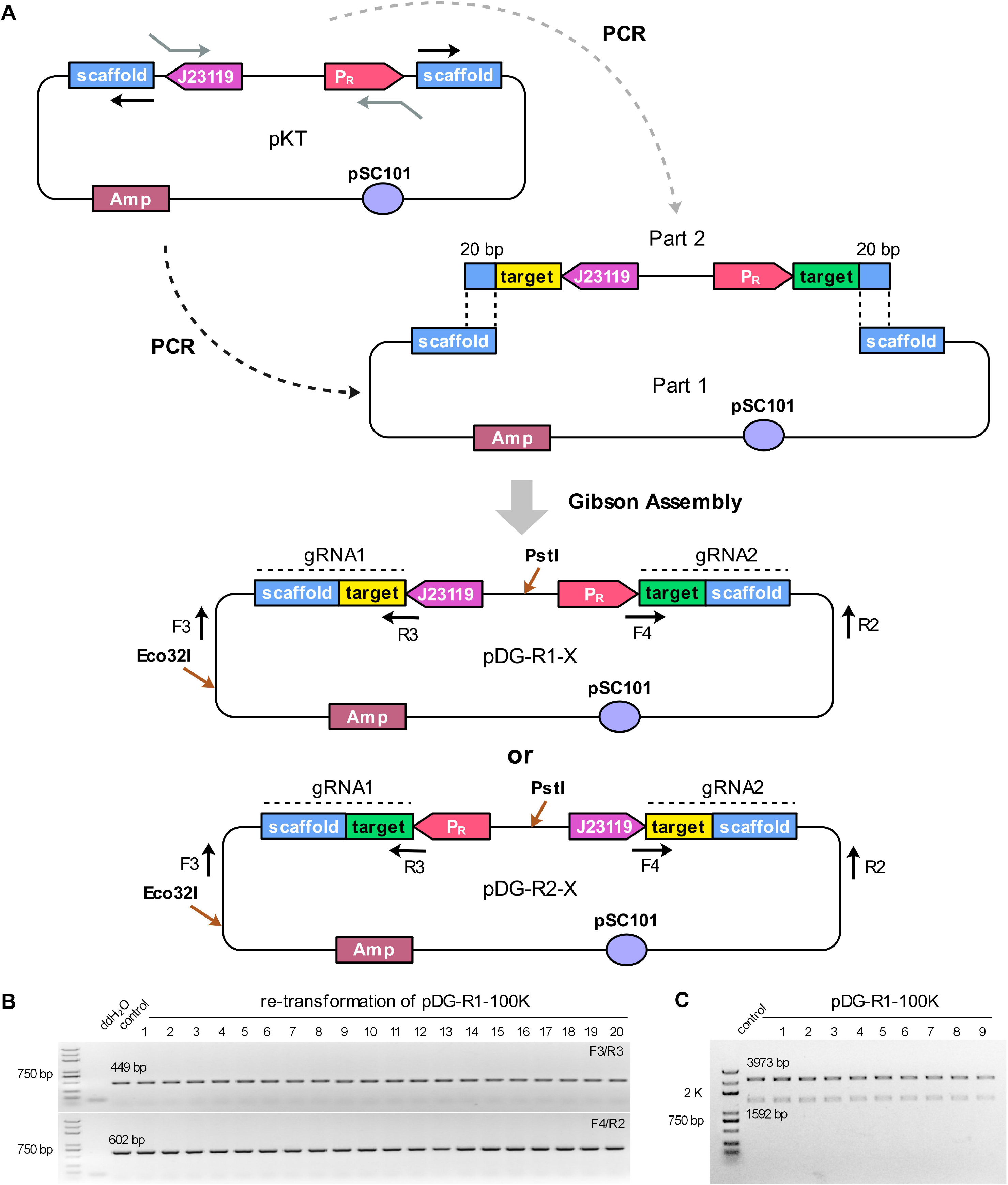
The design and stability of RPGPs pDG-R-100K in *E. coli*. A) The modular construction strategy of pDG-R-X series. pKT plasmid was designed for PCR amplification of DNA part 1 and part 2 series. DNA part 1 which contained pDG-R-X backbone and two reversed repeated gRNA scaffolds was amplified by using only one prime. DNA part 2 contained two different promoters followed by a 20-bp target sequence respectively. For the PCR reaction, the 20-bp sequences specific for two targeted loci and another 20-bp overlap sequences for assembly were embedded in primers as a part of insert. Gibson Assembly method was preformed to assemble these parts into pDG-R1-X or pDG-R2-X series. B) Representative PCR results of pDG-R1-100K after re-transformation process in *E. coli*. C) The double restriction enzyme digestion analyses of pDG-R1-100K and its derivations.

To further examine PRGPs stability, the correct pDG-R1-100K plasmid was retransformed into DH10B strain and verified by PCR reaction. All of 50 colonies produced a 449-bp and a 602-bp band when amplified by primer pair F3/R3 and F4/R2, respectively. Representative colony PCR results are shown in **Figure 5B.** Nine of corresponding plasmids were digested by Eco32I and PstI and produced two bands with correct sizes of 3973 bp and 1592 bp (**Figure 5C**). The following DNA sequencing also confirmed that pDG-R1-100K maintained the intact paired gRNA expression cassettes without any mutations.

### Large genomic deletion mediated by RPGPs

To test the practicability of RPGPs, RPGPs-associated CRISPR/Cas9 system was used for large genome editing in *E. coli* MG1655 strain. Since the double-strand breaks (DSBs) in *E. coli* can be repaired through its native end-joining mechanism (Chaoyong Huang *et al.*, 2019), two plasmids were required for editing: p-P_BAD_-Cas9 plasmid contained the p15A replication origin, a *kan* gene, and Cas9 protein under control of the arabinose-inducible araBAD promoter (P_BAD_); RPGPs pDG-R-K series contained pSC101 replication origin, a *bla* gene and paired-gRNA expression cassettes **(Figure 6A)**. Cas9 used here was an evolved SpCas9 variant xCas9-3.7 (Hu *et al.*, 2018), which could reduce the survival rate of WT cells and increase the positive rate of large genomic editing.

**Figure 6.**
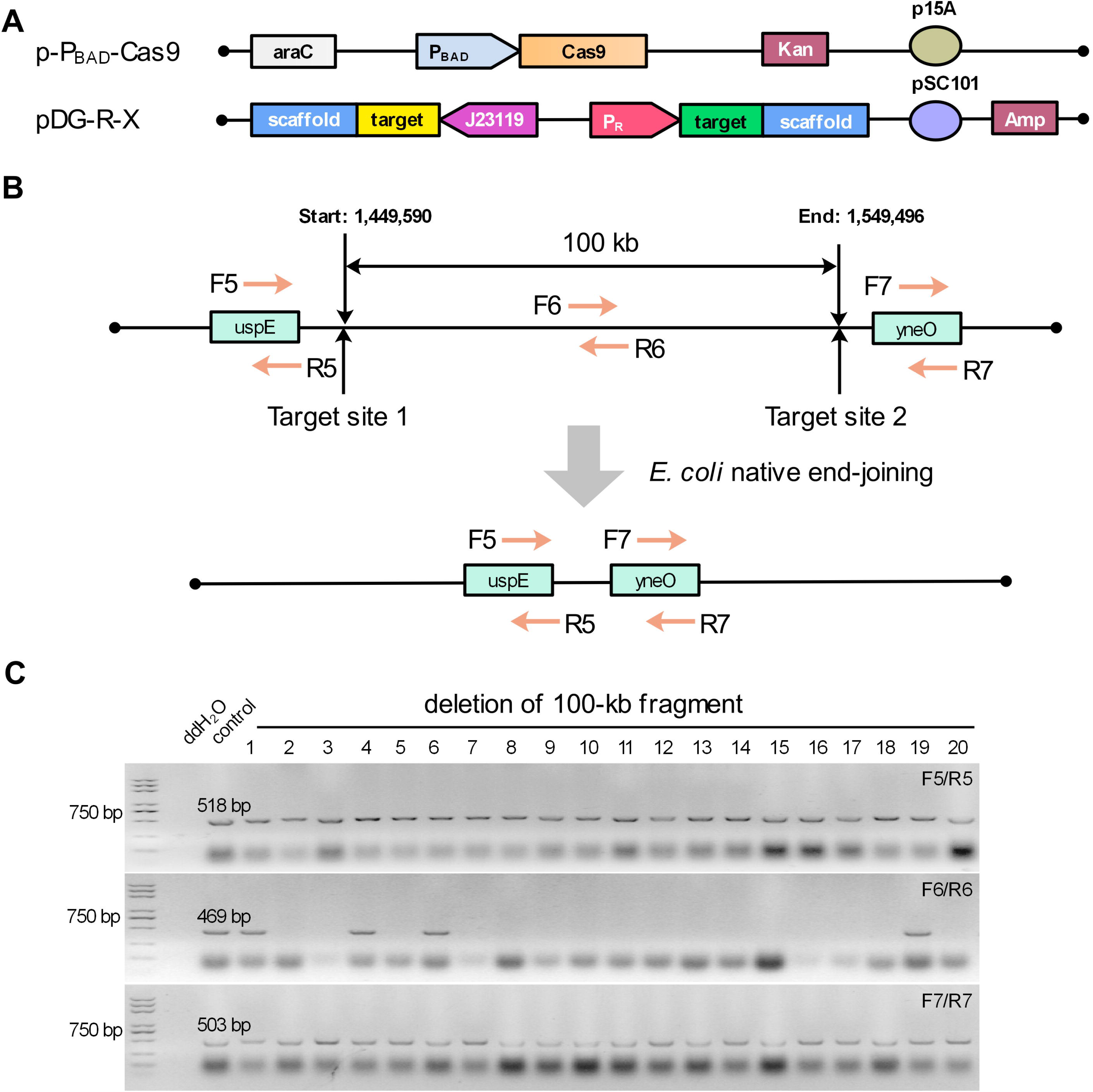
The RPGPs-assisted CRISPR-Cas9 system for the 100-kb fragment deletion in *E. coli*. A) Diagram of p-PBAD-Cas9 and RPGPs pDG-R-X series for genomic deletion. p-PBAD-Cas9 plasmid was used to express Cas9 protein introduced by L-arabinose. pDG-R-X contained two gRNAs for targeting. B) Deletion of a 100-kb genomic fragment in *E. coli*. C) Representative PCR results of a 100-kb fragment deletion. Colonies were randomly picked for PCR screening, and a WT colony served as control.

pDG-R1-100K plasmid was applied to coexpress two gRNAs for the deletion of a 100-kb nonessential fragment from the *E. coli* chromosome (1,449,590-1,549,496) **(Figure 6B)**. The targeting sequences of 100-kb fragment were summarized in **Table S2, Supporting Information**. Three pairs of primer F5/R5, F6/R6, and F7/R7 were designed to check positive mutants among forty randomly selected colonies, and representative PCR results are shown in **Figure 6C**. A proximate 83.33% editing efficiency was achieved in this test, while negative colonies (16.67%) in the experimental group were also obtained. Further investigation showed that these colonies were not the wild type, but contained sequence deletion of stochastic length in the two target sites. The results indicated that RPGPs-associated CRISPR/Cas9 system was successfully used for large genome editing in *E. coli*.

## Discussion

Using *E. coli* as host, this study characterized the rearrangement of plasmids carrying paired gRNAs that appeared as DRs. The plasmid rearrangements were triggered by RecA-independent recombination, leading to spontaneous deletion of one gRNA once introduced into cells. To increase plasmid stability, a variety of modifications toward operating conditions and plasmid architecture were tested by examining the maintenance of paired gRNAs. It showed that electrotransformation of plasmids and the use of nutrient-rich medium increased plasmid stability, but gRNA deletion events could not be eliminated as long as plasmids still contained DRs. Thus, a simple but universal RPGPs cloning strategy was developed by placing paired gRNAs oppositely, or in other words, by transforming DRs into IRs. This strategy made it easy to obtain and maintain the correct plasmids carrying paired gRNAs, and it was efficient in large genomic deletion in *E. coli*.

The deletion mechanism of DRs-involved paired gRNA plasmids was explained with a replication slippage model for RecA-independent recombination (Bi *et al.*, 1995; Saveson and Lovett, 1997). The specific DNA replication process can be predicted as following. When DNA synthesis pauses, the DNA polymerase may dissociate from the DNA sequence, allowing the nascent strand containing the first copy of the DRs to separate from its template strand. Meanwhile, a loop structure formed in between the DRs on the template strand bring the two repeated regions closer, facilitating the nascent strand to translocate and pair to the second copy of the DRs. As a result, the elongation of the nascent strand can skip across the sequence within the looping structure when DNA synthesis resumed, leading to the deletion of one entire copy of DRs (Bzymek and Lovett, 2001), or in this case, one gRNA expression cassette. The point mutations in the promoter and the insertion mutations in the scaffold may be attributed to the interruptions of DNA replication induced by DRs.

Compared to chemical transformation, introducing the gRNA plasmid via electroporation significantly reduced its deletion rate. It is likely that plasmid adsorption by the bacterial membrane will introduce breaks to the DNA strand and result in subsequent removal of the damaged repeats by DNA repair mechanisms, while the plasmids remain intact when entering into cells through the membrane pores via electroporation (Sugar and Neumann., 1984). Also, rearrangements usually occur right after the arrestment of DNA polymerase (Michel and Bénédicte, 2000). The abundant nutrition in the TB medium which accelerates DNA replication and cell division may prevent the nascent strand from translocating and pairing to other repeated regions, thus achieving reduced deletion rate of plasmids.

According to the slipped misalignment model, expansion should be produced as efficiently as deletion (Bzymek and Lovett, 2001). However, the expansion of gRNAs could hardly be observed in this study. This agreed with the single strand annealing model which predicted that only deletion, instead of expansion, was efficiently produced in the presence of DRs separated by palindromes (Bzymek and Lovett, 2001). Although single strand annealing was inefficient in *E. coli* due to the rampant DNA degradation by the RecBCD, it suggested that the secondary structures of the intervening sequence could increase the frequency of deletion. Thus, the hairpin structure within the gRNA scaffold (Jinek *et al.*, 2012) may contribute to the domination of the deletion event in plasmid rearrangement.

The disruption of promoter repeats could increase plasmid stability theoretically, but it was still difficult to obtain the correct plasmid through Gibson Assembly method. Based on the assembly mechanism, we make an assumption that one end of the backbone can be assembled normally to the insert, but another end is mismatched to the repeated gRNA scaffold sequence of insert, due to the help of 5’ exonuclease. Thus, other assembly method *in vitro* or *in vivo* that involved with exonuclease for terminal micro-homologous junction may face the same problem when used for the construction of paired gRNA plasmids. Furthermore, even if the correct plasmid is obtained by certain assembly method, the plasmid stability will be compromised by the two identical gRNA scaffolds during DNA replication. We also found that lowering the copy number of plasmid didn’t increase the plasmid stability as long as DRs excited. In contrast, our RPGPs cloning strategy is simple but useful to avoid DRs-mediated plasmid recombination genetically. Even if IRs-mediated recombination, a very low-frequent recombination, can invert the intervening sequence under some conditions (Bi and Liu, 1996b), it will not jeopardize the expressions of gRNAs in pDG-R-X series.

The genetic basis of the RecA*-*independent recombination is independent of RecBCD, RecET, RecF, RecG, RuvAB and other known recombination mechanisms in *E. coli*, but its mechanism has not been fully understood (Bzymek and Lovett, 2001; Azpiroz and Laviña, 2017). It seems to be difficult to find a suitable host to eliminate RecA*-*independent recombination. Although specific commercial strains (e.g. Stbl3 or NEB stable) with the *recA13* or *recA1* genotype may be able to reduce the frequency of homologous recombination of long repeats, they cannot reduce the occurrence of the RecA-independent recombination. Moreover, even if isolation of plasmid containing repeat elements is facilitated by some strains, the plasmids will again suffer from the high-frequent rearrangement once introduced into the targeted hosts for metabolic engineering. Therefore, our RPGPs strategy which is not limited by host strains have a great potential to be applied for CRISPR-Cas9 systems in various *E. coli* strain with different genotypes.

In summary, the deletion mechanism of DRs-involved paired-gRNA plasmids was investigated and a simple RPGPs cloning strategy for coexpressing paired gRNAs was developed by converting DRs to the more stable IRs. This strategy can completely eliminate DRs-mediated recombination events, improve the performances of CRISPR systems that relied on paired gRNAs, and also facilitate other applications involving repeated genetic parts.

## Experimental procedures

### Strains and Cultivation Conditions

All strains mentioned in this study were listed in **Table S1, Supporting Information**. *E. coli* DH10B and XL10-Gold are *recA1* strains expressing mutated RecA protein which has a G to A point mutation at position of the 482th base for reducing occurrence of nonspecific recombination in cloned DNA. Mach1-T1 Δ*recA1398* strain and DB3.1 Δ(*sr1-recA*) strain have truncated non-functional RecA protein, while MG1655 strain has the intact RecA protein. DH10B was used as a main cloning strain, and MG1655 was used in the genome editing experiments. Luris–Bertani Broth (LB) medium (10 g/L tryptone, 5 g/L yeast extract, 10 g/L NaCl) was used for cell growth in all cases unless otherwise noted. Terrific Broth (TB) medium (12 g/L tryptone, 24 g/L yeast extract, 4 mL/L glycerol, 17mM KH_2_PO_4_:72mM K_2_HPO_4_ buffer solution) was also used for cell growth under some conditions. SOC medium (20 g/L tryptone, 5 g/L yeast extract, 0.5 g/L NaCl, 2.5 mM KCl, 10 mM MgCl2, 10 mM MgSO4, and 20 mM glucose) was used for cell recovery. Twenty g/L agar was supplemented if needed. Antibiotics were added at the following final concentrations: ampicillin, 100 μg/mL; kanamycin, 50 μg/mL. When appropriate, 20 mM L-arabinose were supplemented into media or cultures.

### Plasmids Constructions

The plasmids used in this study were described in **Table S1, Supporting Information**. Target sequences of 100-kb fragment and all primers were given in **Table S2, Supporting Information**. pDG-A-X series, a derivative of pKB, contained ColE1 origin and two identical J23119 promoters (http://parts.igem.org/Part:BBa_J23119) in the same direction to drive paired gRNAs. pDG-P-X series contained a J23119 promoter and a P_R_ promoter (http://parts.igem.org/Part:BBa_R0051) to drive paired gRNAs. pDG-S-X series, a derivative of pKS, contained pSC101 origin and two identical J23119 promoters in the same direction for gRNAs expression. pDG-R-X series, a derivative of pKT, contained pSC101 origin and two different promoters in the opposite direction for co-expression of paired gRNAs. The specific modular construction strategies of pDG-A-X and pDG-R-X series were shown in **Figure 1A** and **Figure 5A**, respectively. All of the paired-gRNA plasmids were constructed by using the Gibson Assembly method which can facilitate two overlapping DNA fragments to be assembled into a circular molecular via the concerted action of a 5’ exonuclease, a DNA polymerase and a DNA ligase (Gibson and Al, 2009). For the PCR reaction, the 20-bp guide sequences specific for two targeted loci were embedded in primers as a part of insert.

All the DNA fragments were PCR-amplified with Phusion polymerase (New England BioLabs). PCR products were gel purified, digested with *Dpn*I enzyme (Thermo Fisher Scientific) before assembly, which could eliminate the template plasmids. Gibson Assembly^®^ Master Mix were ordered from New England BioLabs.

### Deletion assays

When the plasmids or their assembly products were introduced into *E. coli* strains, twenty colonies on the Ap^r^ plate were selected randomly and verified by colony PCR in each experiment. The deletion rate was determined by the ratio of the number of mutated colonies among twenty randomly-picked colonies. Three biological replicates were tested for each group of experiments unless otherwise noted.

### Genome editing procedure

*E. coli* MG1655 containing p-P_BAD_-cas9 (Chaoyong Huang *et al.*, 2019) electrocompetent cells were prepared firstly. Then, specific plasmid pDG-R1-100K (100 ng) were added in each electroporation reaction. Bio-Rad MicroPulser was used for electroporation (0.1 cm cuvette, 1.80 kV). Cells after electroporation were immediately added into 1 mL SOC medium and recovered for 45 min prior to plating on LB (Ap^r^+ Kan^r^) plate at 30 °C. A single colony was picked and inoculated into 0.5 mL SOC medium and cultured at 30 °C for 2 h. Then, 4.5 mL LB (Ap^r^+ Kan^r^) medium were added to the cultures. After 1 h, L-arabinose (20 mM) was added, and the cultures were cultured for another 3 h before plating. A 10-μL aliquot of the cultures was plated onto a LB (Ap^r^+ Kan^r^) plate containing L-arabinose, overnight at 30 °C. To analyze the mutation types of colonies, PCR products of the target locus were cloned and subjected to DNA sequencing.

## Supporting Information

Supporting Information is available from the Wiley Online Library or from the author.

## Acknowledgements

This study was jointly supported by the National Natural Science Foundation of China (grant No. 21676026), the National Key R&D Program of China (grant No. 2017YFD0201400) and Fundamental Research Funds for the Central Universities.

## Reference

Aparicioprat, E., Arnan, C., Sala, I., Bosch, N., Guigó, R., and Johnson, R. (2015) DECKO: Single-oligo, dual-CRISPR deletion of genomic elements including long non-coding RNAs, J Bmc Genomics 17: 215.

Azpiroz, M.F., and Laviña, M. (2017) Analysis of RecA-independent recombination events between short direct repeats related to a genomic island and to a plasmid in *Escherichia coli* K12, PeerJ 5: e3293.

Bi, X., and Liu, L.F. (1994) recA-independent and recA-dependent intramolecular plasmid recombination. Differential homology requirement and distance effect, J Mol Bio 235: 414.

Bi, X., Lyu, Y.L., and Liu, L.F. (1995) Specific stimulation of recA-independent plasmid recombination by a DNA sequence at a distance, J Mol Bio 247: 890–902.

Bi, X., and Liu, L.F. (1996a) A replicational model for DNA recombination between direct repeats, J Mol Bio 256: 849–858.

Bi, X., and Liu, L.F. (1996b) DNA rearrangement mediated by inverted repeats, Proc Natl Acad Sci USA 93: 819–823.

Bzymek, M., and Lovett, S.T. (2001) Instability of repetitive DNA sequences: the role of replication in multiple mechanisms, Proc Natl Acad Sci USA 98: 8319–8325.

Chaoyong Huang, Tingting Ding, Jingge Wang, Xueqin Wang, Liwei Guo, Jialei Wang et al. (2019) CRISPR-Cas9-assisted native end-joining editing offers a simple strategy for efficient genetic engineering in *Escherichia coli*, Appl Microbiol Biotechnol 103: 8497–8509.

Cox, M.M., and Lehman, I.R. (1987) Enzymes of General Recombination, Annu Rev Biochem 56: 229–262.

Del, S., G., Giraldo, R., Ruiz-Echevarría, M.J., Espinosa, M., and Díaz-Orejas, R. (1998) Replication and control of circular bacterial plasmids, Microbiol Mol Biol R62: 434.

Gibson, D.G., and Al, E. (2009) Enzymatic assembly of DNA molecules up to several hundred kilobases, Nat Methods 6: 343.

Hasunuma, K., and Sekiguchi, M. (1977) Replication of plasmid pSC101 in *Escherichia coli* K12: requirement for dnaA function, Mol Gen Genet 154: 225–230.

Hu, J.H., Miller, S.M., Geurts, M.H., Tang, W., Chen, L., Sun, N. et al. (2018) Evolved Cas9 variants with broad PAM compatibility and high DNA specificity, Nature 556.

Jiang, W., Bikard, D., Cox, D., Zhang, F., and Marraffini, L.A. (2013) RNA-guided editing of bacterial genomes using CRISPR-Cas systems, Nat Biotechnol 31: 233–239.

Jiang, Y., Chen, B., Duan, C., Sun, B., Yang, J., and Yang, S. (2015) Multigene editing in the *Escherichia coli* genome via the CRISPR-Cas9 System, Appl Environ Microbiol 81: 2506.

Jinek, M., Chylinski, K., Fonfara, I., Hauer, M., Doudna, J.A., and Charpentier, E. (2012) A programmable dual-RNA-guided DNA endonuclease in adaptive bacterial immunity, Science 337: 816.

Kowalczykowski, S.C. (1991) Biochemical and biological function of *Escherichia coli* RecA protein: behavior of mutant RecA proteins, Biochimie 13: 289–304.

Li, Y., Lin, Z., Huang, C., Zhang, Y., Wang, Z., Tang, Y.J. et al. (2015) Metabolic engineering of Escherichia coli using CRISPR–Cas9 meditated genome editing, Metab Eng 31: 13.

Lovett, S.T., Drapkin, P.T., Jr, S.V., and Gluckmanpeskind, T.J. (1993) A sister-strand exchange mechanism for recA-independent deletion of repeated DNA sequences in *Escherichia coli*, Genetics 135: 631–642.

Lovett, S.T., and Feschenko, V.V. (1996) Stabilization of diverged tandem repeats by mismatch repair: evidence for deletion formation via a misaligned replication intermediate, Proc Natl Acad Sci USA 93: 7120–7124.

Radding, C.M. (1989) Helical RecA nucleoprotein filaments mediate homologous pairing and strand exchange, Biochimica et biophysica acta 1008: 131.

Ran, F.A., Hsu, P., Lin, C.Y., Gootenberg, J., Konermann, S., Trevino, A.E. et al. (2013) Double Nicking by RNA-Guided CRISPR Cas9 for Enhanced Genome Editing Specificity, Cell 154: 1380–1389.

Reis, A.C., Halper, S.M., Vezeau, G.E., Cetnar, D.P., Hossain, A., Clauer, P.R., and Salis, H.M. (2019) Simultaneous repression of multiple bacterial genes using nonrepetitive extra-long sgRNA arrays, Nat Biotechnol 31: 1294–1301

Saveson, C.J., and Lovett, S.T. (1997) Enhanced deletion formation by aberrant DNA replication in *Escherichia coli*, Genetics 146: 457–470.

Su, T., Liu, F., Gu, P., Jin, H., Chang, Y., Wang, Q. et al. (2016) A CRISPR-Cas9 Assisted Non-Homologous End-Joining Strategy for One-step Engineering of Bacterial Genome, Sci Rep 6: 37895.

Sugar, I.P., and Neumann., E. (1984) Stochastic model for electric field-induced membrane pores electroporation, Biophys Chem 19: 211–225.

Vidigal, J.A., and Ventura, A. (2015) Rapid and efficient one-step generation of paired gRNA CRISPR-Cas9 libraries, Nat Commun 6: 8083.

Zheng, X., Li, S.Y., Zhao, G.P., and Wang, J. (2017) An efficient system for deletion of large DNA fragments in *Escherichia coli* via introduction of both Cas9 and the non-homologous end joining system from *Mycobacterium smegmatis*, Biochem Bioph Res Co 485: 768.

