## Supplementary material for "Reversed paired-gRNA plasmid cloning strategy for efficient genome editing in *Escherichia coli*": Table S1, Supporting Information

### Supporting Data

### Tables

**Table S1.** Strains and plasmids used in this study.

| Strains or plasmids | Descriptions | Source |
|---|---|---|
| **Strains** | | |
| *E. coli* DH10B | *$F^-$ $\lambda^-$ endA1 recA1 mcrA galU galK nupG rpsL ΔlacX74 Δ(mrr-hsdRMS-mcrBC) φ80lacZΔM15 araDΔ139 Δ(ara-leu)7697 ($str^r$)* | Invitrogen |
| *E. coli* XL10-Gold | *endA1 recA1 thi-1 gyrA96 relA1 lac The Δ(mcrA)183 Δ(mcrCB-hsdSMR-mrr)173 $tet^r$ F'[proAB $lacI^qZ$ΔM15 Tn10($Tet^r$ Amy $Cm^r$ ]* | Invitrogen |
| *E. coli* Mach1-T1 | *$F^-$ endA1 ΔrecA1398 tonA φ80(lacZ)ΔM15 ΔlacX74 hsdR($rk^-$ $mk^+$ )* | Invitrogen |
| *E. coli* DB3.1 | *$F^-$ endA1 Δ(sr1-recA) gyrA462 mcrB mrr hsdS20(rB-,mB-) supE44 ara-14 galK2 lacY1 proA2 rpsL20 xyl-5λ-leumtl1* | Biomed |
| *E. coli* MG1655 | *$F^-$ $\lambda^-$ ilvG- rfb-50 rph-1* | Our lab |
| **Plasmids** | | |
| pKB | ColE1; J23119 promoter, a gRNA scaffold; $Ap^r$ | This study |
| pKI | ColE1; J23119 promoter, a gRNA scaffold; $Ap^r$ | This study |
| pPR | ColE1; $P_R$ promoter, a gRNA scaffold; $Ap^r$ | This study |
| pKS | pSC101; J23119 promoter, a gRNA scaffold; $Ap^r$ | This study |
| pKT | pSC101; J23119 promoter, $P_R$ promoter, two gRNA scaffolds; $Ap^r$ | This study |
| pDG-A-100K | ColE1; two J23119 promoters, two gRNAs; $Ap^r$ | This study |
| pDG-P-100K | ColE1; J23119 promoter, $P_R$ promoter, two gRNAs; $Ap^r$ | This study |
| pDG-S-100K | pSC101; two J23119 promoters, two gRNAs; $Ap^r$ | This study |
| pDG-R-100K | pSC101; J23119 promoter, $P_R$ promoter, two gRNAs; $Ap^r$ | This study |
| p-$P_{BAD}$-cas9 | p15A; *Cas9*; $Kan^r$ | [1] |

**Table S2.** Target sequences of 100-kb fragment and all primers used in this study.

**Target sequences of 100-kb fragment.**

| Spacer | Nucleotide sequence (5'-3') | PAM |
|---|---|---|
| 100K-1 | gtaaaaccgaatactgccgg | tgg |
| 100K-2 | gaagtggctaaagagaacaa | cgg |

**Primers for construction of CPGPs and RPGPs.**

| Primer Name | Primer sequence (5'-3') |
|---|---|
| pKB backbone _F | gttttagagctagaaatagc |
| pKB backbone_R | actagtattatacctaggac |
| pDG-A-100K insert_F | gtcctaggtataatactagtgtaaaaccgaatactgccggttttagagctagaaatagc |
| pDG-A-100K insert_R | gctatttctagctctaaaacttgttctctttagccacttcactagtattatacctaggac |
| pDG-P-100K insert_F | gtcctaggtataatactagtgtaaaaccgaatactgccggttttagagctagaaatagc |
| pDG-P-100K insert_R | ctatttctagctctaaaacttgttctctttagccacttcgcaaccattatcaccgcca |
| pKS backbone _F | gttttagagctagaaatagc |
| pKS backbone_R | actagtattatacctaggac |
| pDG-S-100K insert_F | gtcctaggtataatactagtgtaaaaccgaatactgccggttttagagctagaaatagc |
| pDG-S-100K insert_R | gctatttctagctctaaaacttgttctctttagccacttcactagtattatacctaggac |
| pKT backbone_F/R | gttttagagctagaaatagc |
| pDG-R-100K insert_F | tatttctagctctaaaacccggcagtattcggttttacactagtattatacctagg |
| pDG-R-100K insert_R | tatttctagctctaaaacttgttctctttagccacttcgcaaccattatcaccgcc |

To clone pDG-A-100K, primers pKB backbone _F/pKB backbone _ R were used to amplify pKB, and primers pDG-A-100K insert_F/pDG-A-100K insert_R were used to amplify pKI. The PCR products were assembled through Gibson Assembly method [2].

To clone pDG-P-100K, primers pKB backbone _F/pKB backbone _ R were used to amplify pKB, and primers pDG-P-100K insert_F/pDG-P-100K insert_R were used to amplify pPR. The PCR products were assembled through Gibson Assembly method [2].

To clone pDG-S-100K, primers pKS backbone _F/pKS backbone _ R were used to amplify pKS, and primers pDG-S-100K insert_F/pDG-S-100K insert_R were used to amplify pKI. The PCR products were assembled through Gibson Assembly method [2].

To clone pDG-R-100K, primers pKT backbone _F/pKT backbone _ R were used to amplify pKT, and primers pDG-R-100K insert_F/pDG-R-100K insert_R were used to amplify pKT. The PCR products were assembled through Gibson Assembly method [2].

**Screening and sequencing primers for CPGPs and RPGPs.**

| Primer Name | Primer sequence (5'-3') |
|---|---|
| F1 | ttgacagctagctcagtcct |
| R1 | ctctgctaatcctgttaccag |
| F2 | taacaccgtgcgtgttgact |
| R2 | ttggtggttgataagcgagg |
| F3 | gatcactacttcgcactagt |
| R3 | taatactagtgtaaaaccga |
| F4 | taatggttgcgaagtggcta |
| gRNA-Sequencing_1 | aacgcggcctttttacggttc |
| gRNA-Sequencing_2 | tccccgaaaagtgccacctg |
| gRNA-Sequencing_3 | actacacgatgctttaactg |

For screening of pDG-A-100K, primers F1/R1 were used. Primer gRNA-Sequencing_1 was used for DNA sequencing.
For screening of pDG-P-100K, primers F1/R1 or F2/R1 were used. Primer gRNA-Sequencing_1 was used for DNA sequencing.
For screening of pDG-S-100K, primers F1/R2 were used. Primer gRNA-Sequencing_2 was used for DNA sequencing.
For screening of pDG-R1-100K, primers F3/R3 and F4/R2 were used respectively. Primers gRNA-Sequencing_2 and gRNA-Sequencing_3 were used for DNA sequencing.

**Screening and sequencing primers for 100-kb genome editing.**

| Primer Name | Primer sequence (5'-3') |
|---|---|
| F5 | gcaaatgttgcccacgaatg |
| R5 | ggacgaacgtaccgctatg |
| F6 | cggtcgatatgcggatgtat |
| R6 | tctcctgctgcaggattttg |
| F7 | gacaacaagcccctgattac |
| R7 | gaactggggaggcgactatt |

### Reference

[1] Chaoyong Huang, Tingting Ding, Jingge Wang, Xueqin Wang, Liwei Guo, Jialei Wang, Lin Zhu, Changhao Bi, Xueli Zhang, Xiaoyan Ma, Y.-X. Huo, *Applied Microbiology and Biotechnology* **2019**, *103*, 8497–8509.
[2] D. G. Gibson, E. Al, *Nature Methods* **2009**, *6*, 343.
